# RepliCNN: High-resolution inference of the DNA replication program from strand-specific 3′ DNA end sequencing

**DOI:** 10.64898/2026.03.12.710907

**Authors:** Dominik Stroh, Nicola Zilio, Maruthi K. Pabba, Vassilis Roukos, M. Cristina Cardoso, Helle D. Ulrich, Kathi Zarnack

**Affiliations:** Theodor Boveri Institute, University of Würzburg, 97074 Würzburg, Germany; Institute of Molecular Biology (IMB) gGmbH, 55128 Mainz, Germany; Technical University of Darmstadt, 64287 Darmstadt, Germany; Department of Biology, Medical School, University of Patras, Patras, Greece; Cluster for Nucleic Acid Sciences and Technologies – NUCLEATE

**Keywords:** Replication timing, origin of replication, Okazaki fragments, CNN, strand-specific 3′ DNA end sequencing, TrAEL-seq, OK-Seq, GLOE-Seq

## Abstract

During S phase, the genome is replicated in a tightly regulated spatiotemporal order described as DNA replication timing (RT). Discontinuous lagging-strand synthesis produces Okazaki fragments whose strand-specific distribution reflects replication dynamics. Here, we present RepliCNN, a deep learning framework based on one-dimensional convolutional neural networks to predict RT from Okazaki fragment distributions obtained from strand-specific 3′ DNA end sequencing methods such as GLOE-Seq, TrAEL-seq, or OK-Seq. RepliCNN also automatically annotates replication origins, termination zones, replication fork directionality, and origin efficiency genome-wide from a single dataset. Benchmarking on public and in-house human and yeast datasets using leave-one-chromosome-out cross-validation demonstrates high predictive accuracy in both wild-type and perturbation experiments, enabling comprehensive analyses of replication dynamics from strand-specific DNA 3′ end sequencing data.

**Highlights:**

- RepliCNN enables integrated analysis of replication timing, fork directionality and replication features from strand-specific 3′ DNA end sequencing data.
- High-resolution replication dynamics can be inferred from a single experiment, bypassing complex multi-fraction labelling approaches.
- The framework generalizes across experimental protocols, datasets, and species.
- This enables cost-effective comparative analysis of replication programs across biological conditions.

## Background

In eukaryotic cells, each chromosome must replicate precisely once per cell cycle prior to mitosis. Any perturbations of this process may result in the loss of genetic information due to incomplete genome duplication or copy number alterations, thereby compromising genome stability and promoting disease, including various forms of cancer (Gaillard et al., 2015). Accordingly, DNA replication is governed by multilayered regulatory mechanisms that coordinate origin licensing, origin firing, and fork progression in both space and time.

During S phase, replication initiates at specific genomic loci termed origins of replication. The structure and composition of origins differ substantially across eukaryotes. In the budding yeast *Saccharomyces cerevisiae*, origins are well-defined elements characterized by conserved sequence motifs and precisely mapped binding sites for the origin recognition complex and associated factors. In contrast, in metazoans, initiation events occur within broad initiation zones rather than at sharply defined single sites. Upon origin activation, the replicative helicase unwinds the parental duplex, generating a replication bubble from which two replication forks emanate bidirectionally. At each fork, DNA synthesis proceeds in a semi-discontinuous manner: the leading strand is synthesized continuously in the direction of fork movement, whereas the lagging strand is synthesized discontinuously as short Okazaki fragments that are subsequently processed and ligated. Genome-wide replication profiles can be partitioned into distinct domain domains: Constant timing regions comprise segments of the genome that undergo replication synchronously within a defined window of S phase. These domains are interconnected by timing transition regions, which reflect extended unidirectional replication fork progression emanating from neighboring early activated regions. Replication termination regions correspond to broad regions where oppositely moving replication forks converge and DNA synthesis is completed.

Although simultaneous activation of all licensed origins might, in principle, maximize replication efficiency, only a subset of origins fires in any given S phase, and their activation follows a defined temporal order referred to as replication timing (RT). RT correlates strongly with transcriptional output and epigenetic landscape, thereby coordinating replication with gene expression (Vouzas and Gilbert, 2023). Consequently, RT profiles are reprogrammed during embryogenesis and cell fate transitions (Hiratani et al., 2008; Nakatani et al., 2024) and commonly exhibit aberrations in cancer and other disease contexts (Dietzen et al., 2024). Mechanistically, it has been postulated that origins compete for activation factors or nucleotides, resulting early firing of the most efficient origins (Mantiero et al., 2011). An alternative, non-mutually exclusive hypothesis suggests that dormant origins serve as a backup that can be activated in response to replication stress, thereby mitigating the impact of stalled replication forks and safeguarding genome integrity (Blow and Ge, 2009). Several genomic features distinguish early replicating regions which are typically gene-rich, highly transcribed, enriched for open chromatin marks (Vouzas and Gilbert, 2023; González-Acosta and Lopes, 2024), and preferentially located in the nuclear center, often in proximity to nuclear speckles (Chen et al., 2018). In contrast, late replicating regions are characterized by a paucity of genes, low levels of transcription, heterochromatin, and proximity to the nuclear lamina or the nucleolus (Bersaglieri et al., 2022). It has also been proposed that essential or highly expressed genes replicate early to allow extended time for DNA damage detection and repair or because their accessible chromatin environment facilitates early origin activation (Stamatoyannopoulos et al., 2009; Dietzen et al., 2024). In higher eukaryotes, RT is strongly linked to three-dimensional genome organization, as the timing of DNA replication is typically established domain-wise at the level of DNA loops or topologically associating domains during G1 phase (Pope et al., 2014; Dileep et al., 2015).

The spatial and temporal orchestration of DNA replication is crucial for cellular function, yet analyzing these processes genome-wide remains a bottleneck. The state-of-the-art approaches for RT profiling in cells, including Repli-Seq, involve bromodeoxyuridine (BrdU) incorporation into actively replicating DNA, followed by cell sorting of S-phase fractions, BrdU immunoprecipitation, high-throughput sequencing, and bioinformatics analyses (Hansen et al., 2010). High-resolution implementations using 16 fractions provide near-continuous temporal information across S phase (Zhao et al., 2020). Other variants of this approach have been developed to optimize cost efficiency (Marchal et al., 2018), allow for single-cell profiling (Takahashi et al., 2019), or bypass cell sorting and immunoprecipitation using nanopore-based strategies (Theulot et al., 2025). While these methodologies allow to profile replication, they are labor-intensive, require substantial cell numbers, extensive sequencing depth, and complex experimental workflows, making them costly and limiting scalability. Moreover, they are inherently limited in their capacity to precisely map origins. These can be more accurately resolved by dedicated strand-specific 3′ DNA end sequencing protocols that aim to directly capture of Okazaki fragments (OK-Seq) (Smith and Whitehouse, 2012; Petryk et al., 2016), unligated 3′ (OH) ends (GLOE-Seq, TrAEL-seq) (Sriramachandran et al., 2020; Kara et al., 2021), replication bubbles (Bubble-Seq) (Mesner et al., 2013), short nascent DNA strands (SNS-seq) (Gómez and Brockdorff, 2004), or initiation sites (INI-seq) (Langley et al., 2016; Guilbaud et al., 2022).

In stark contrast to the overwhelming number of experimental protocols available for RT profiling and origin identification, the computational landscape is sparse. Existing tools address specific aspects of replication analysis, such as RT prediction from primary DNA sequence, processing of single-cell replication timing data (Gnan et al., 2022), general RT characterization (Brison et al., 2019), analysis and visualization of OK-Seq data (Liu et al., 2023), or prediction of replication fork directionality (Arbona et al., 2023). However, these tools are typically designed for narrowly defined applications and tailored to a specific experiment type.

Here, we present RepliCNN, a deep learning-based, integrated analysis framework to overcome these limitations. RepliCNN allows to predict RT and to identify origins, initiation zones, and termination zones for diverse types of strand-specific 3′ DNA end sequencing data. Importantly, RepliCNN obviates the need for labor-intensive, multi-fraction experiments by extracting high-resolution RT information directly from single-fraction datasets such as TrAEL-seq or related methods. This substantially reduces experimental complexity, cost, and time requirements, while enabling systematic comparison across biological conditions. In this study, we demonstrate the cross-applicability of RepliCNN between samples, experiment types and species.

## Results

### RepliCNN is a framework to analyze replication dynamics from strand-specific 3′ DNA end sequencing data

High-resolution Repli-Seq data provide detailed spatiotemporal information on RT and enable the delineation of replication domains, yet such multi-fraction experiments are labor-intensive, costly, and experimentally demanding. Conceptually, much of the information required to reconstruct replication dynamics is already encoded in strand-specific 3′ DNA end sequencing data. Approaches such as GLOE-Seq, TrAEL-seq, or OK-Seq generate genome-wide, strand-resolved profiles of nascent DNA fragments, including Okazaki fragments or unligated 3′ hydroxyl ends. Because lagging-strand synthesis produces discontinuous fragments, asymmetric accumulation of open 3′ ends on the Watson or Crick strand reflects the predominant direction of replication fork movement within a genomic region. Transitions in strand bias occur at sites of replication initiation and termination, thereby marking origins and termination zones. This principle is exemplarily illustrated using TrAEL-seq data from wild-type *S. cerevisiae* cells (**Figure 1**, top left), highlighting how strand asymmetry inherently encodes spatiotemporal features of DNA replication.

**Figure 1.**
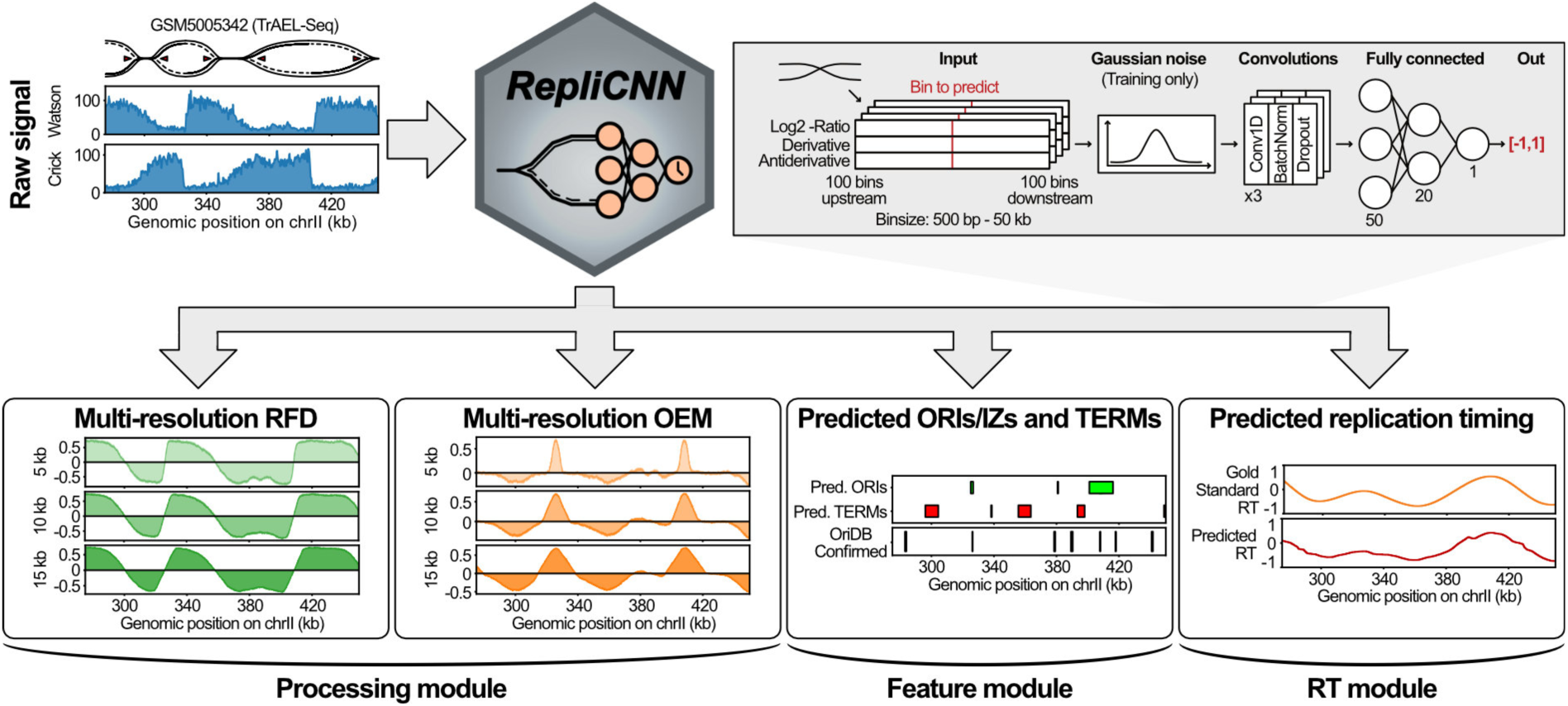
RepliCNN workflow for analyzing DNA replication dynamics from strand-specific 3′ DNA end sequencing data. RepliCNN takes raw strand-specific 3′ DNA end sequencing data generated by protocols such as GLOE-Seq, TrAEL-seq, or OK-Seq as input. The workflow comprises three integrated modules: (i) The data processing module computes multi-resolution replication fork directionality (RFD) and origin efficiency metric (OEM) tracks using variable window sizes and strides. (ii) The feature module identifies origins of replication (ORIs) and termination zones (TERMs) and estimates their genomic widths based on the derived signal tracks. (iii) The RT module comprises a convolutional neural network that predicts RT profiles following training on orthogonal data such as Repli-Seq. Once trained on a wild-type profile, the model can be transferred to other datasets.

To leverage the inherent spatiotemporal RT information, we developed RepliCNN, a computational framework for the analysis of strand-specific 3′ DNA end fragment sequencing datasets, including GLOE-Seq, TrAEL-seq, OK-Seq, and related methodologies. RepliCNN exploits the inherent strand asymmetry to quantitatively infer replication fork directionality and origin activity, thereby enabling high-resolution mapping of RT domains including origins and termination zones. RepliCNN thereby allows to reconstruct DNA replication dynamics across multiple genomic scales.

The RepliCNN workflow comprises three integrated components (**Figure 1**). First, the data processing module computes multi-resolution replication fork directionality (RFD) and origin efficiency metric (OEM) (McGuffee et al., 2013) tracks using user-defined window sizes and strides. RFD is derived from strand-specific read densities and reflects the relative abundance of rightward- versus leftward-moving replication forks in a given window. OEM is calculated from local changes in RFD and provides a quantitative measure of origin firing efficiency. By generating these signals across multiple spatial resolutions, RepliCNN captures both sharp, focal initiation events and broader replication domains without relying on excessive smoothing or fixed binning strategies.

Second, the feature module identifies candidate origins (ORIs) and termination zones (TERMs) from the derived signal tracks. This module implements a multi-step procedure that (i) detects candidate initiation and termination sites at each resolution, (ii) integrates calls across resolutions, (iii) recenters and resizes candidate sites based on local signal maxima or inflection points, and (iv) applies quantitative thresholds to define high-confidence ORIs and TERMs. By combining information across resolutions, this approach increases robustness and reproducibility relative to single-resolution or heavily smoothed methods, which may obscure narrow origins or inflate broad domains.

Third, RepliCNN incorporates a one-dimensional convolutional neural network designed to predict RT directly from the strand-specific 3′ DNA end profiles (RT module; **Figure 1**, top right). During training, the model learns to map processed strand asymmetry features to orthogonal RT datasets (e.g., Repli-Seq) derived from matched wild-type samples. Once trained, the model can be applied to perturbed or other experimental conditions lacking direct RT measurements, thereby enabling inference of genome-wide RT shifts.

For model input, genomic data are segmented into overlapping windows comprising 201 consecutive bins, with the central bin representing the prediction target. In budding yeast, bins typically span 500 bp, whereas in larger genomes (e.g., human) bin sizes range from 10–50 kb. From each window, RepliCNN computes a log₂ ratio of Watson and Crick strand signals, analogous to RFD, and additional features including spline-based approximations as well as their first derivative and integral. These transformations capture both local fork polarity and broader replication domain structure.

During training, feature tensors are regularized via a Gaussian noise layer and passed through three one-dimensional convolutional blocks, each consisting of a convolutional layer followed by batch normalization and dropout. The resulting representations are flattened and processed by a fully connected network comprising three dense layers (50, 20, and 1 neuron, respectively). A hyperbolic tangent activation function constrains the final output to the interval [–1, 1], corresponding to late (negative) and early (positive) RT states. This architecture enables accurate and transferable prediction of RT from strand-specific 3′ DNA sequencing data alone.

RepliCNN can be downloaded from GitHub and installed via pip or ran as an Apptainer (former Singularity) container. It does not require expert knowledge in bioinformatics, can be efficiently integrated into pipelines and used as an importable package for python scripts and notebooks. RepliCNN also provides data loaders and savers for all supported data types. RepliCNN’s all-in-one solution thus provides a unified framework to analyze strand-specific 3′ DNA end datasets and predict RT and other features in a reproducible manner.

### RepliCNN accurately predicts replication timing and origins of replication in yeast

To validate our approach, we applied RepliCNN to strand-specific 3′ DNA end sequencing data from *Saccharomyces cerevisiae*, whose compact genome makes it a well-suited model organism for studying DNA replication (Bell and Labib, 2016). We first evaluated the performance of the RepliCNN RT module using TrAEL-seq and OK-Seq data from wild-type yeast cells (Smith and Whitehouse, 2012; Kara et al., 2021) as well as GLOE-Seq data from *Δcdc9* (DNA ligase I) cells which accumulate unligated Okazaki fragment ends (Sriramachandran et al., 2020; Kara et al., 2021). Using the strand-specific signal, we used the RepliCNN data processing module to calculate replication fork directionality (RFD) and origin efficiency metric (OEM) tracks at three resolutions (5 kb, 10 kb, and 15 kb window size). Gold-standard RT information was derived from (Müller and Nieduszynski, 2012) and smoothed to attenuate local outliers prior to model training (see Materials and Methods). Models were trained using a sample-wise leave-one-chromosome-out cross-validation (LOCO-CV) scheme, whereby a model was trained for each target chromosome with that chromosome excluded from the training data, precluding data leakage. Predictive accuracy was quantified using Pearson correlation coefficients (PCC). We observed strong agreement between predicted and gold-standard RT, with chromosome-wise median correlation coefficients exceeding 0.8 (**Figure 2A**), demonstrating the robustness of RepliCNN’s RT predictions across diverse experimental modalities.

**Figure 2.**
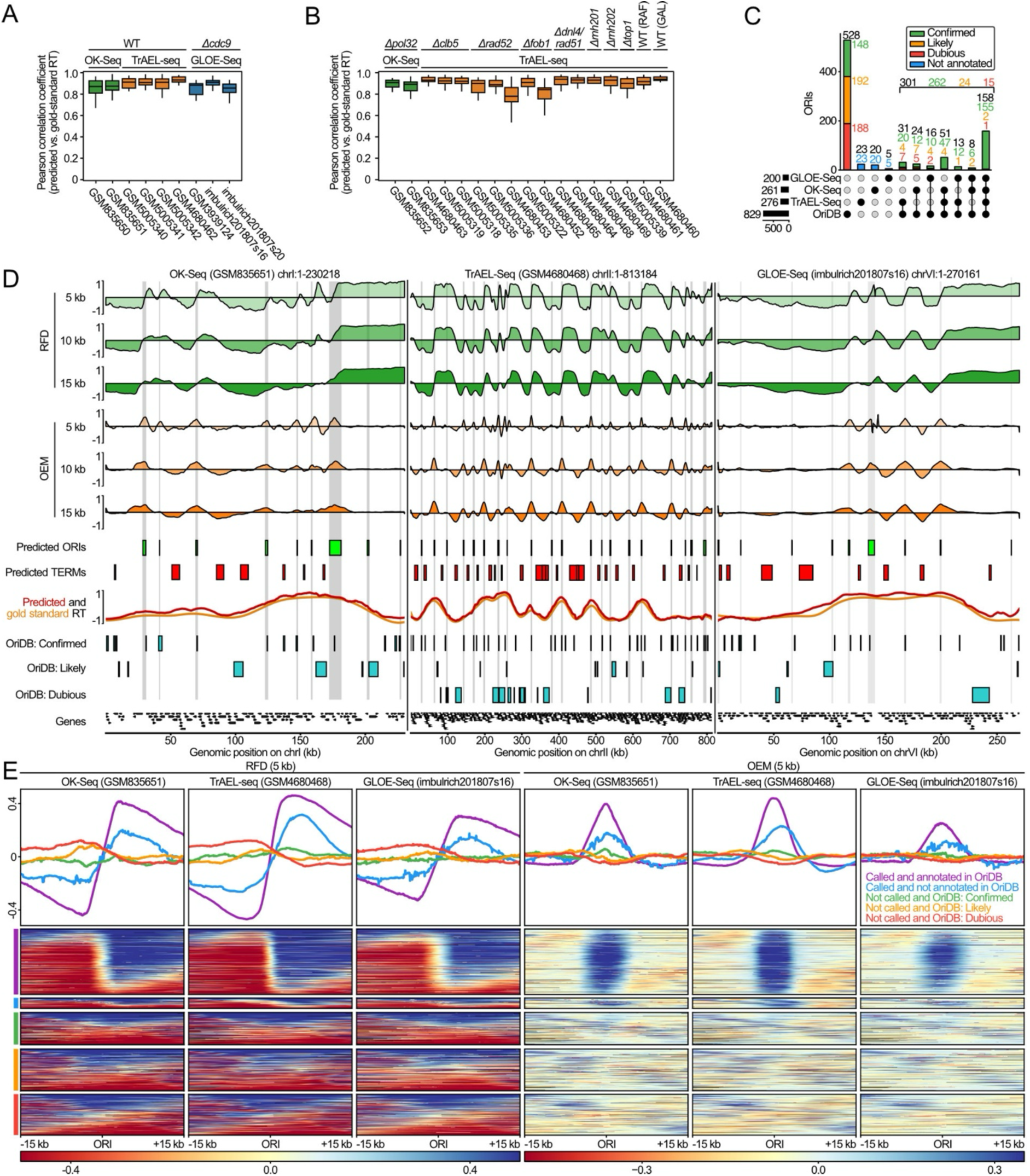
RepliCNN performance and origin calling for strand-specific 3′ DNA end sequencing data from *S. cerevisiae*. **(A)** Barplot showing Pearson correlation coefficients between predicted and gold-standard RT for yeast OK-Seq, TrAEL-seq, and GLOE-Seq samples collected from multiple studies. GLOE-Seq data for *Δcdc9* (DNA ligase I) cells were used for comparison as these accumulate unligated Okazaki fragment ends. **(B)** Barplot as in (A) for OK-Seq and TrAEL-seq data for various yeast mutant strains and growth conditions (raffinose [RAF] or galactose [GAL] as sugar source) as indicated. **(C)** UpSet plot of predicted and annotated origins of replication (ORIs, retrieved from OriDB) for three representative samples from each experiment type. Overlaps between unannotated ORIs across experiments are not shown. **(D)** Representative regions for each experiment type showing multi-resolution replication fork directionality (RFD) and origin efficiency metric (OEM) tracks. Predicted ORIs and termination zones (TERMs), as well as predicted and gold-standard RT, inferred from (Müller and Nieduszynski, 2012), are shown below. Also shown are OriDB origins of the three categories (confirmed, likely, and dubious) and genes overlapping with each region. **(E)** RFD and OEM profiles and heatmaps around ORIs for the three experiment types, stratified by the category determined in (B).

We further extended the analysis to the full set of publicly available datasets for mutant yeast strains from the three experimental protocols, including e.g. OK-Seq for *Δpol32* cells (polymerase δ subunit, impaired strand-displacement processivity) (Smith and Whitehouse, 2012) or TrAEL-seq data for *Δclb5* cells (cyclin B, activation of late-firing origins) (Kara et al., 2021). As previously reported, the RT profiles in the mutant strains were generally similar to wild-type across the genome, reflected in a high correlation of the RepliCNN-predicted RT profiles to the gold-standard from wild-type (**Figure 2B**). Some samples showed more divergent patterns, for instance for *Δrad52* cells (DNA repair & homologous recombination) or *Δfob1* cells (unidirectional rDNA replication), albeit with high variability between replicates.

We next assessed the capacity of the RepliCNN feature module to identify origins of replication by benchmarking predicted ORIs against annotations from the yeast OriDB database (Smith and Whitehouse, 2012). Using a representative sample for each experiment type, we identified 276 ORIs from TrAEL-seq, 200 ORIs from GLOE-Seq, and 261 ORIs from OK-Seq (**Figure 2C, D**). Notably, the majority of RepliCNN-predicted ORIs were shared across sequencing modalities, with only a minor fraction of experiment-specific ORIs identified (**Figure 2C**). Quantitatively, 156 OriDB-annotated ORIs were recovered across all three experiment types. Among the 528 OriDB-annotated ORIs that remained undetected across all experimental modalities, 380 (72%) were flagged as “likely” or “dubious” annotations within OriDB.

To further substantiate that RepliCNN predicts *bona-fide* origins, we evaluated the aggregated RFD and OEM signal at the predicted sites. ORIs detected by both RepliCNN and OriDB displayed the most pronounced origin signature, i.e. a sign switch from negative to positive RFD values and a corresponding OEM peak—indicative of high-confidence active origins—in all three experiments (**Figure 2E**). Importantly, ORIs predicted by RepliCNN but absent from OriDB also exhibited an intermediate yet clearly discernible signature, suggesting that RepliCNN recovers a set of bona-fide active origins lacking in current OriDB annotations. Conversely, OriDB-annotated ORIs not identified by RepliCNN showed no detectable RFD sign switch, suggesting that these origins may be dormant or only rarely fire under the experimental conditions. The minor OEM enrichment visible for these sites may reflect low-efficiency origins that are frequently passively replicated by forks originating from neighboring origins. Under such conditions, forks do not consistently emanate in both directions, yet local changes in fork polarity can still give rise to a detectable OEM signal.

Taken together, these results demonstrate that RepliCNN accurately reconstructs both RT profiles and origin maps from strand-specific 3′ DNA end sequencing data in budding yeast. The strong agreement with gold-standard RT, the consistent detection of known origins across experimental protocols, and the identification of additional sites with clear origin-like signatures collectively highlight the robustness and sensitivity of the framework.

### RepliCNN captures initiation zones and replication timing in human cells

Having established the performance for budding yeast, we next investigated whether RepliCNN can similarly capture replication dynamics in more complex eukaryotic genomes. To test this, we performed TrAEL-seq for the human colorectal carcinoma cell line HCT116 (*n* = 3 replicates). As gold-standard RT information, we used a high-resolution Repli-Seq dataset (16 fractions) for the same cell line (Zhao et al., 2020). Using RepliCNN, we first computed multi-resolution RFD and OEM tracks (50 kb, 75 kb, 100 kb, and 150 kb window size) and assessed their reproducibility across replicates. RFD tracks showed consistently high inter-replicate correlation, uniformly exceeding 0.90 (**Figure 3A**). PCCs increasing up to 0.97 at the coarsest resolution reflecting that larger window sizes attenuate local noise and yield more reproducible directional estimates (Petryk et al., 2016). OEM tracks exhibited comparatively lower inter-replicate correlation, yet with a similar resolution-dependent increase, indicating a slightly greater inherent variability in efficiency relative to directionality estimates.

**Figure 3.**
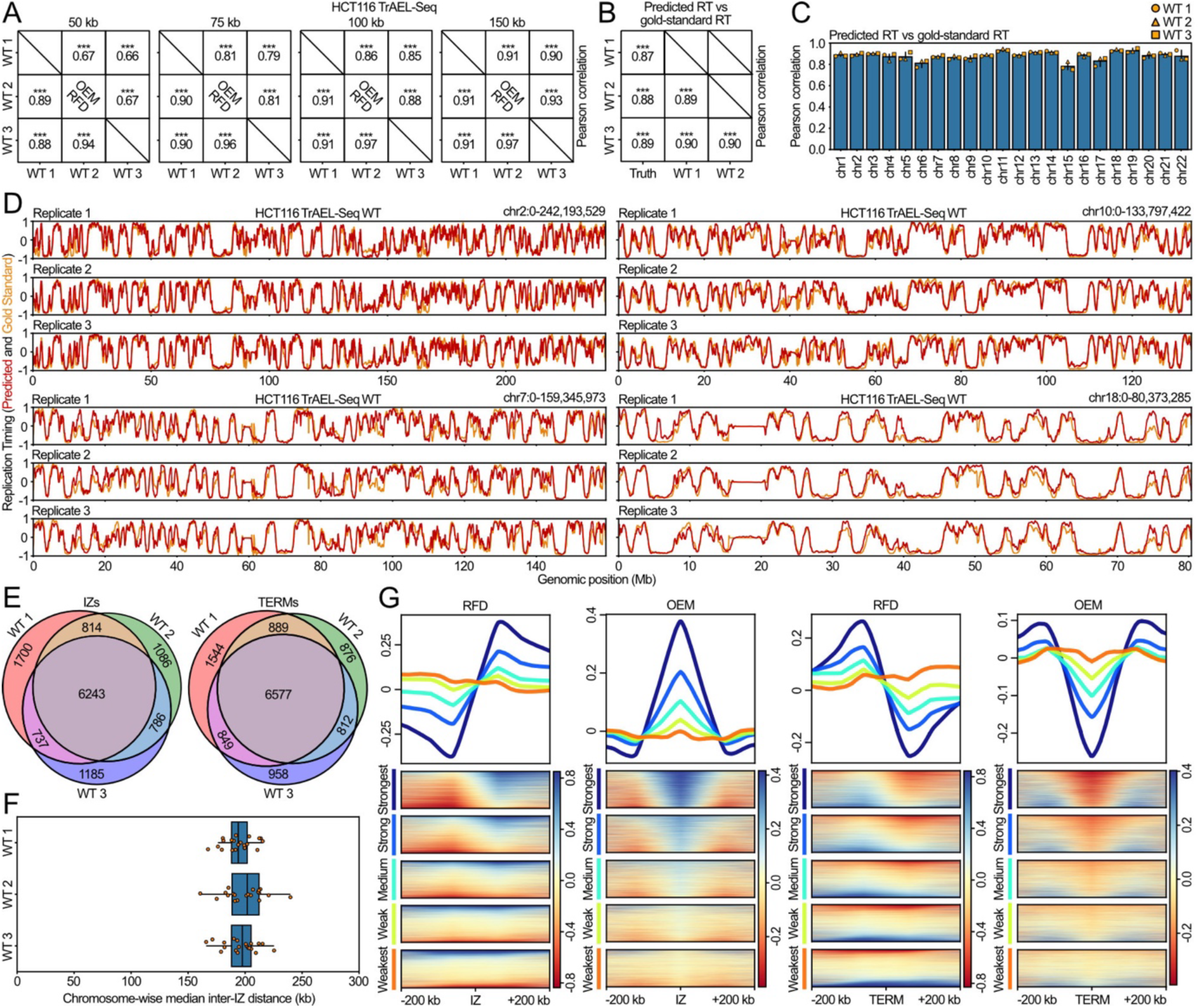
RepliCNN accurately predicts human replication timing and discovers replication initiation zones. **(A)** Pearson correlation coefficient heatmaps of RFD (lower triangular) and OEM (upper triangular) for four window sizes (50 kb, 75 kb, 100 kb, and 150 kb; three leftmost panels) and predicted versus gold-standard RT from (Zhao et al., 2020) (rightmost panel), derived from a human HCT116 TrAEL-seq wildtype experiment. Statistical significance is indicated by asterisks: *p<0.05, **p<0.01, ***p<0.001. **(B)** Barplot of chromosome-wise Pearson correlation coefficients between predicted and true (gold standard) replication timing. **(C)** Predicted versus gold-standard RT for four representative chromosomes across all three HCT116 TrAEL-seq wildtype replicates. Values of 1 and −1 indicate the earliest and latest replication timing, respectively. **(D)** Venn diagrams showing the overlap between predicted initiation zones (IZs) and termination zones (TERMs) across the three replicates. IZs and TERMs were considered overlapping if located within 50 kb of each other. **(E)** Boxplot of chromosome-wise median inter-IZ distances. Outliers were identified using the interquartile range method and are not shown. **(F)** Profiles and heatmaps of RFD and OEM signal around IZs and TERMs. IZs and TERMs are stratified by signal strength (see Materials and Methods for details) into quintiles, where 80–100 represents the strongest and 0–20 the weakest IZs and TERMs.

We subsequently applied the RepliCNN RT module to each replicate using a LOCO-CV scheme. The predicted RT profiles exhibited high agreement with the gold-standard across all samples, with PCCs > 0.85 (**Figure 3B**), demonstrating that RepliCNN’s predictive capacity extends to the considerably larger and more complex human genome. Representative examples from chromosomes 2, 7, 10, and 18 illustrate the strong concordance between RepliCNN predictions and gold-standard RT profiles across the three replicates (**Figure 3C**).

Using the RepliCNN feature module, we identified origins and termination zones from the multi-resolution RFD and OEM tracks. Following the terminology in metazoans—and matching the increased window size compared to the yeast data analysis—we refer to the origins in the human genome as initiation zones (IZs). Across replicates, a total of 12,551 IZs were identified, of which 8,580 (68%) were reproducibly detected in at least two replicates (**Figure 3D**). Analogously, 9,127 out of a total of 12,505 (73%) TERMs were found in at least two replicates, supporting the robustness of RepliCNN’s feature identification (**Figure 3D**). The identified IZs showed a median inter-IZ distance of approximately 200 kb (**Figure 3E**), consistent with previously reported inter-origin distances in mammalian cells (Cayrou et al., 2011; Petryk et al., 2016). Stratifying the identified IZs by feature strength, we observed that the 20% strongest IZs exhibited the most pronounced RFD sign switch (from negative to positive), accompanied by the sharpest OEM peak, while progressively weaker tiers displayed attenuated signatures (**Figure 3F**). However, even the lowest 20% still displayed a mild peak in the OEM signal, possibly reflecting low-efficiency origins (see above). TERMs displayed a reciprocal pattern, with the strongest 20% exhibiting the most pronounced sign switch from positive to negative RFD values and correspondingly deeper OEM troughs.

Together, these results demonstrate that RepliCNN accurately reconstructs replication timing and replication features from strand-specific 3′ DNA end sequencing data in the human genome, despite its substantially larger size and increased regulatory complexity. This highlights the general applicability of RepliCNN as a scalable framework for extracting high-resolution replication dynamics from single-condition datasets.

### RepliCNN models generalize across datasets, experimental modalities, and species

We next investigated how the choice of training sample influences predictive performance across datasets, experimental modalities, and species. To this end, we assembled a compendium of 32 samples, comprising 22 TrAEL-seq (19 for yeast, 3 for human) (Kara et al., 2021), 6 GLOE-Seq (3 for yeast, 3 for human) (Sriramachandran et al., 2020), 4 OK-Seq (all yeast) experiments (Smith and Whitehouse, 2012).

To comprehensively evaluate generalizability, we defined two training regimes. In the intra-species regime, models were trained and evaluated using an adapted LOCO-CV scheme in which all but one chromosome from the training sample were used for model fitting, and the held-out chromosome subsequently predicted in the target sample. This setup enabled comparisons both within and between experimental protocols for the same species while preventing information leakage between samples. For cross-species evaluation—which precludes data leakage by design—models were trained on the complete dataset of the training organism.

Across the evaluation panel, samples displayed a noticeable variability in performance, both as training data and as prediction targets, indicating differences in data quality and/or biological variability among experiments (**Figure 4**). Predictive performance was highest within experiments, likely reflecting the ability of each model to exploit sample-specific characteristics in its training data (**Figure 4A**). For instance, models trained on GLOE-Seq or TrAEL-seq data from human HCT116 cells demonstrated strong inter-replicate predictability (regions ① and ②). Notably, the models retained their predictive power across experimental protocols; for instance, models trained on yeast OK-Seq data performed comparably well when applied to yeast OK-Seq and TrAEL-seq data (regions ③ and ④). An intriguing asymmetry was observed in the cross-species applications. While models trained on human datasets yielded robust predictions for yeast samples, the reciprocal approach—training on yeast data and predicting human samples—resulted in substantially lower performance (regions ⑤ and ⑥). More broadly, while most samples fell below the diagonal in the training-versus-prediction performance comparison, yeast samples were notable exceptions, exhibiting only moderate utility as training data yet high predictability as prediction targets (**Figure 4B–D**).

**Figure 4.**
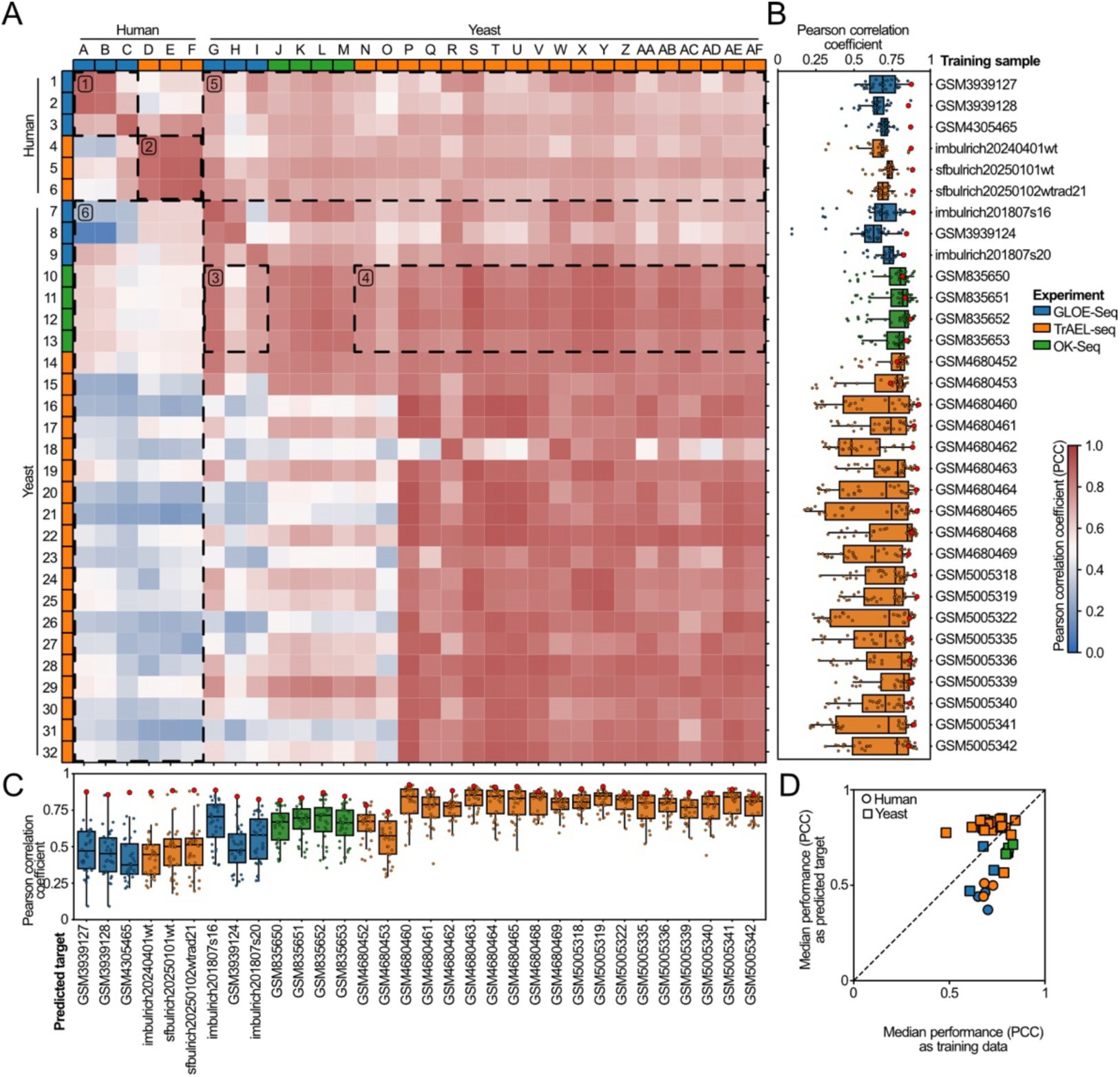
RepliCNN models are applicable across datasets, protocols, and species. **(A)** Heatmap of Pearson correlation coefficients between predicted and gold-standard RT. Rows indicate the sample on which each model was trained; columns indicate the target sample being predicted. A modified leave-one-chromosome-out cross-validation (LOCO-CV) approach was used to avoid information leakage for intra-species comparisons. Dashed rectangles and encircled numbers highlight predictive power between replicates (① GLOE-Seq, ② TrAEL-seq ②, yeast), between experimental protocols (OK-seq-trained model applied to GLOE-Seq ③ and TrAEL-seq ④), and between species (human-trained model applied to yeast ⑤, and vice versa ⑥). **(B,C)** Summarized performances for each sample as training data (B) or prediction target (C). Boxplot of row-wise (B) or column-wise (C) Pearson correlation coefficients per sample from heatmap in (A). Red dots indicate prediction performance within the same sample (i.e., value on the diagonal). **(D)** Scatterplot of median performance (Pearson correlation coefficient) as training data (x-axis) or prediction target (y-axis) for each sample. Shapes and colors indicate organism and experimental protocol, respectively.

Taken together, these results demonstrate that RepliCNN shows high performance in inter-experiment settings, where a model trained on a reference RT profile can be used to infer RT in related or perturbed conditions. The observed cross-species asymmetry suggests that models trained on the more complex replication landscapes of larger genomes capture generalizable features that remain informative in simpler systems such as yeast.

## Discussion

Genome-wide replication profiling methods such as BrdU-Seq, Repli-Seq, OK-Seq, GLOE-Seq, and TrAEL-seq provide complementary views of replication dynamics by encoding information on fork directionality, origin positioning and replication timing. However, the computational infrastructure supporting these assays has remained fragmented, typically consisting of single-purpose tools tailored to specific experimental modalities. RepliCNN addresses this gap by providing a unified computational framework that infers RT and replication features from strand-specific 3′ DNA end sequencing data. Annotation of origins and termination zones is performed using rule-based methods without requiring assay-specific parameter retuning. By learning multi-scale representations of strand asymmetry across configurable genomic windows, the framework reduces the reliance on fixed-bandwidth smoothing that can obscure narrow origins or artificially inflate replication domains, while remaining applicable across multiple experimental protocols and species.

A central observation of this study is that strand-specific 3′ DNA end distributions contain sufficient information to accurately reconstruct genome-wide replication timing. Previous work demonstrated that strand-specific Okazaki fragment polarity reflects replication fork directionality and switches at replication origins, indicating that such data captures key aspects of replication dynamics (McGuffee et al., 2013; Petryk et al., 2016; Sriramachandran et al., 2020; Kara et al., 2021). Our results extend this concept by showing that these signals encode enough information to infer the complete genome-wide RT profile. While RT has traditionally been measured using multi-fraction labeling experiments such as high-resolution Repli-Seq, we demonstrate that comparable information can be extracted from single-condition datasets such as TrAEL-seq. This substantially lowers the experimental costs for RT analysis and facilitates comparative studies across perturbations or biological conditions in which dedicated assays may not be feasible.

The robust predictive performance of RepliCNN across two evolutionarily distant organisms indicates that the features learned by the convolutional neural network capture general properties of replication dynamics rather than species-specific signatures. Leave-one-chromosome-out validation provides a conservative estimate of generalization performance by preventing leakage of chromosomally correlated features during training. We show that the framework performs consistently for the defined origin architecture of *S. cerevisiae* as well as the broad initiation zones characteristic of mammalian genomes. Compatibility across multiple sequencing modalities further supports the robustness of the approach, since each method introduces distinct biases in fragment recovery, genomic coverage and strand asymmetry patterns.

Traditional pipelines rely on fixed bin sizes combined with heavy smoothing procedures such as Gaussian or Loess fitting. Although these approaches reduce noise, they can suppress fine-scale origin signals and introduce biases due to manually defined thresholds. RepliCNN’s convolutional architecture instead learns to weight genomic scales according to their predictive value. This allows the model to capture both the sharp local asymmetries associated with discrete origins and the broader gradients that define timing domains. Although downstream feature annotation in RepliCNN still relies on defined thresholds, the data-driven RT inference likely reduces sensitivity to noise in individual bins and improves robustness across datasets.

Taken together, RepliCNN offers an integrated computational framework for analyzing replication patterns from strand-specific 3′ DNA end distributions derived from TrAEL-seq, GLOE-Seq, OK-Seq, and related experimental protocols. By integrating multi-resolution signal processing with deep learning-based inference, RepliCNN enables comprehensive characterization of DNA replication dynamics from a single experimental modality. As experimental technologies for mapping DNA replication continue to diversify, computational approaches capable of extracting shared biological signals across experimental modalities will become increasingly important for decoding the spatiotemporal organization of genome replication.

## Material and methods

### Cell culture

All cells used were tested for mycoplasma and deemed free of contamination. All cell lines were authenticated by STR profiling (ATCC). Human colorectal carcinoma cells (HCT116, https://www.atcc.org/products/ccl-247) were cultured in RPMI-1640 (R8758, Sigma, Germany) supplemented with 10% fetal bovine serum (FBS-12A, Capricorn scientific GmbH, Germany), 1 mM sodium pyruvate (S8636, Sigma Aldrich Chemie GmbH, Germany), 1× L-glutamine (392-0441, VWR, Germany) and 50 µg/ml gentamicin (A005, HiMedia, India) at 37° C with 5% CO_2_. Once the cells reached ∼70% confluency, the medium was removed, and the cells were washed 1× with PBS. Next, the cells were dissociated using 0.25% trypsin-EDTA (TRY-3B, Capricorn GmbH, Germany) at 37° C, followed by inactivation with culture medium. The cells were then pelleted at 300 ×*g* for 5 min, washed once with 1× PBS and pelleted again.

### Agarose plug preparation

To prepare agarose-cell plugs, 2% low-melting-point agarose in water (50081, Lonza, Switzerland) was used. Per plug, 1.5–2 million cells were used. All other steps were carried out as described in (Kara et al., 2021) and https://www.babraham.ac.uk/sites/default/files/2024-11/Embedding-cells-in-agarose.doc.

### TrAEL-seq library preparation

TrAEL-seq libraries were produced as described in (Kara et al., 2021) and https://www.babraham.ac.uk/sites/default/files/2022-03/TrAEL-seq.doc, with a few minor changes. (i) After the initial ligation with T4 RNA ligase 2 KQ, the plugs were transferred to 2 ml tubes and washed 8× (∼ 1 h each) with 2 ml 1× Tris buffer at room temperature before leaving them overnight with 2 ml 1× Tris buffer on a rocker. (ii) After agarose digestion, the DNA was captured with 50 µl SPRI beads (HighPrep PCR, AC-60050, MagBio, USA) + 200 µL spared SPRI bead buffer, washed twice with 80% ethanol and resuspended in 25 µl water. The DNA, together with the SPRI beads, was carried over to the next extension step with Bst 2.0 polymerase. (iii) After this extension step, the DNA was captured on the SPRI beads by supplementing the reaction with 50% PEG-8000 and 5 M NaCl to a final concentration of 7.5% and 0.9 M, respectively, washing with 80% ethanol and eluting in 130 µl 1× TE. (iv) Washed DNA captured onto MyOne streptavidin C1 was resuspended in 25 µl water and transferred to 8-tube PCR strips for the subsequent end polishing reaction. All subsequent steps, until the end of the procedure, were carried out in these tubes. Any intervening washes were accordingly reduced to 200 µl, instead of 500 µl, as described in the original procedure. In addition, for these steps, 1× IDTE (11-01-02-05, IDT, USA) was used instead of 1× TE and Thermolabile USER^®^ II Enzyme (M5508, New England Biolabs, USA).

### High-throughput sequencing

Libraries were sequenced on an Illumina NextSeq 2000 sequencer with P1/P2/P3-100 flow cells, based on how many libraries were sequenced and how many reads were required at a time. All libraries were sequenced in paired-end mode, with read lengths of 2×50 bases plus 6+6 or 8+8 bases for the indices. Sequencing depth was between 40 and 60 million reads per sample. Upon completion of the run, raw sequencing reads of pooled libraries were demultiplexed based on their index sequences by BCLconvert (Illumina). The resulting FASTQ files were used as input for subsequent analysis.

### Processing of published strand-specific 3′ DNA end sequencing data

Published TrAEL-seq data for yeast were downloaded from NCBI Gene Expression Omnibus (GEO) with accession number GSE154811. Published and newly generated data was processed as follows: Read preprocessing was conducted as described in the original publication. In short, the unique molecular identifiers (UMIs) were removed from each read and written into its header. Up to three poly-thymins (polyT) were trimmed from the 5′ end of the read. After TrimGalore (https://github.com/FelixKrueger/TrimGalore) was run with default parameters. Reads were then mapped to the reference genome (human genome version hg38 or yest genome version sacCer3) with bowtie2 (Langmead and Salzberg, 2012) and standard options except for --local. Afterwards, UmiBam (https://github.com/FelixKrueger/Umi-Grinder) was used for UMI-based read deduplication. Strand-specific genome-wide read counts were then generated with bedtools genomecov (Quinlan and Hall, 2010) with options -bg -strand [+/-] −5 and converted to BIGWIG files using kentutils bedGraphToBigWig (https://github.com/ucscGenomeBrowser/kent).

Published human and yeast GLOE-Seq data were downloaded from NCBI GEO with accession number GSE134225. We added two more samples that had been part of the original set of experiments described in (Sriramachandran et al., 2020). The data was processed using GLOE-Pipe (Petrosino et al., 2020) with default parameters as described using genome versions sacCer3 for yeast and hg38 for human.

Published processed OK-Seq data were downloaded from NCBI GEO with accession number GSE33786. BED files were lifted over from genome version sacCer2 to sacCer3 using kentutils liftOver (https://github.com/ucscGenomeBrowser/kent). BED files were then split per strand, truncated at the 5′ end with bedtools genomecov (Quinlan and Hall, 2010), and converted to BIGWIG files using kentutils bedGraphToBigWig (https://github.com/ucscGenomeBrowser/kent).

### Replication timing processing

Repli-Seq data for wild-type human HCT116 cells and replication timing data for wild-type yeast cells were downloaded from NCBI GEO with accession numbers GSE137764 and GSE36045, respectively. Yeast data was lifted from sacCer1 to sacCer3 using kentutils liftOver (https://github.com/ucscGenomeBrowser/kent), and annotated centromeric regions were removed from human data. Afterwards, replication timing was approximated using a SciPy (Virtanen et al., 2020) univariate spline (human: no smoothing, yeast: smoothing factor s = length of chromosome/100_000). The splines were then evaluated in fixed bin sizes (human: 50 kb, yeast: 500 bp), resulting in one BEDGRAPH file with timing associated to each genomic bin for each organism. Timing values were scaled into the interval 1 (early) to –1 (late).

### Identification of origins of replication and termination zones

First, the processed strand-specific 3′ DNA end sequencing were used to compute multi-resolution replication fork directionality (RFD) and origin efficiency metric (OEM) tracks.

Multi-resolution RFD was calculated with the formular:

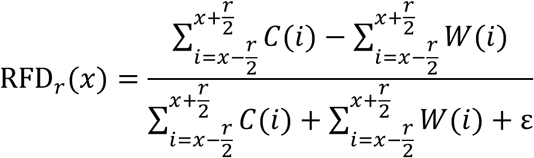

where x is the genomic position, r the resolution, and ε a small number to avoid zero-division errors, and C(i) and W(i) give the signal at position *i* on the Crick and Whatson strand, respectively, for each genomic bin given by the user-specified stride s.

The multi-resolution OEM was calculated with the formular:

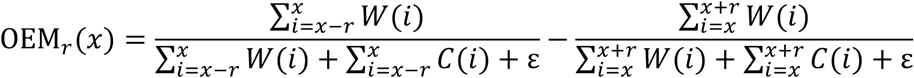

with variables as above.

To systematically identify ORIs and TERMs from the RFD OEM tracks, we developed a multi-step computational pipeline described in detail below.

Step 1 – Candidate identification: Candidate ORIs and TERMs are independently identified at each specified resolution using two complementary approaches. First, RFD signal tracks in bigWig format are smoothed using a univariate spline (SciPy (Virtanen et al., 2020); smoothing factor user-defined) and analytically evaluated to detect zero-crossings: transitions from negative to positive RFD values are designated as candidate ORI positions, whereas transitions from positive to negative RFD values are designated as candidate TERM positions. Second, OEM signal tracks at each resolution are likewise approximated by a univariate spline, and local maxima and minima of the resulting spline are extracted as candidate ORIs and TERMs, respectively. This dual-evidence strategy ensures that candidates are supported by both fork directionality and Okazaki fragment asymmetry signals.

Step 2 – Candidate merging: Candidate ORIs and TERMs identified across all resolutions were subsequently merged into consensus clusters. Candidates located within a user-defined distance threshold of one another were grouped into a single cluster, independently for ORIs and TERMs. This step consolidates spatially proximate calls arising from different resolutions or signal sources into unified genomic intervals.

Step 3 – Resolution-based filtering: Each consensus cluster was then filtered based on the number of independent resolution-level calls supporting it. Specifically, clusters were retained only if they were supported by a minimum number of RFD- and/or OEM-derived candidates across resolutions, as determined by a user-specified evidence threshold. This step eliminates spurious candidates that lack cross-resolution reproducibility.

Step 4 – Recentering and resizing: To refine the precise location and extent of each candidate region, clusters were recentered and resized using OEM data at a user-specified resolution. For each cluster, the local OEM extremum nearest to the cluster center was identified within a user-defined maximum search window; the position of this extremum was used to recenter the cluster. Cluster boundaries were subsequently adjusted based on the amplitude of the local OEM extremum (i.e., peak height for ORIs or trough depth for TERMs) relative to a user-defined threshold, such that the final interval reflects the genomic extent of enriched OEM signal.

Step 5 – Quantification of candidate strength: The strength of each candidate ORI and TERM was quantified using OEM signal data at a user-specified resolution. The strength metric was defined as the maximum OEM signal within a candidate ORI interval or the minimum OEM signal within a candidate TERM interval, providing a measure of the initiation or termination activity at each locus.

Step 6 – Final filtering by signal cutoff: In a final filtering step, candidate ORIs and TERMs were retained only if their quantified strength exceeded a user-defined signal cutoff. This step removes low-confidence calls with weak OEM support, yielding a high-confidence set of replication origins, initiation zones, and termination zones for downstream analyses.

### Training and application of RepliCNN

#### RepliCNN Prepare

Strand-specific sequencing signal tracks in bigWig format, representing forward and reverse strand read densities, were processed to compute log₂-ratio signals. The genomes were partitioned into non-overlapping bins of user-defined size, and mean per-bin strand coverage was extracted using the average_over_bed function from the pybigtools library (Huey and Abdennur, 2024). The log₂-ratio was computed as the log₂ ratio of forward to reverse strand coverage, with a pseudocount (ε = 10⁻¹⁵) added to avoid undefined values in zero-coverage regions. To reduce high-frequency noise while preserving large-scale directionality patterns, a cubic interpolating spline (SciPy UnivariateSpline, *k* = 3) was independently fitted to the ratio signal of each chromosome. The first derivative and antiderivative of each spline were analytically derived, and all three tracks were z-score normalized per chromosome. Optionally, all signal tracks could be inverted to accommodate differing strand labeling conventions, and replication timing data in bedGraph format could be incorporated by mapping timing values onto the bin coordinates. The resulting features, the per-bin strand coverage, log₂ RFD, spline-smoothed RFD, its derivative, antiderivative, and replication timing, were assembled into a standardized data matrix serving as input for all downstream analyses.

#### RepliCNN Train

Input features were constructed from the standardized data matrix by extracting three per-bin signals—log₂ RFD, its first derivative, and its antiderivative—alongside replication timing values as regression targets. For each genomic bin, a sliding window of user-defined size (default: 201 bins) was centered on the bin of interest, with zero-padding applied at chromosome boundaries, yielding a three-channel input tensor of shape (window size × 3 features) per sample. The convolutional neural network (CNN) architecture is implemented using PyTorch (Paszke et al., 2019) and keras (https://keras.io) and consisted of an input Gaussian noise layer (σ = 0.1) for regularization, followed by three successive convolution–batch normalization–dropout blocks with 64, 32, and 16 filters and kernel sizes of 32, 16, and 8, respectively, all using ReLU activations and a dropout rate of 0.5. The convolutional output was flattened and passed through two fully connected layers (50 and 20 units, ReLU) before a single output neuron with a tanh activation produced the replication timing prediction. The model was compiled with the Adam optimizer (Kingma and Ba, 2014) (default learning rate: 0.001) and mean squared error loss, and trained for up to 300 epochs with a batch size of 1,024. Early stopping was applied by monitoring training loss with a minimum improvement threshold of 10⁻³ and a patience of 20 epochs, restoring the best-performing weights upon termination. A user-defined fraction of training data (default: 10%) was held out as a validation set during training. To assess generalization performance, an optional leave-one-chromosome-out cross-validation (LOCO-CV) routine was implemented, in which the model was iteratively trained on all chromosomes except one and evaluated on the held-out chromosome. Predictive performance for each held-out chromosome was assessed by computing the Pearson correlation coefficient between predicted and observed replication timing values. All random operations were seeded for reproducibility (seed = 42), and training was performed on either CPU or GPU depending on hardware availability.

#### RepliCNN Predict

Replication timing was predicted genome-wide by applying the trained CNN model to the standardized data matrix of a query sample. Input features were constructed from the query data using the same sliding window procedure described above, yielding a feature tensor of identical shape to that used during training. The trained model was then applied to this feature tensor to generate per-bin replication timing predictions, which were rounded to three decimal places and written back into the standardized data matrix, replacing or populating the replication timing column. As during training, inference was performed on either CPU or GPU depending on hardware availability, and all random operations were seeded for reproducibility (seed = 42).

#### Software requirements

RepliCNN is written in Python 3 and depends on NumPy (Harris et al., 2020), pandas (https://doi.org/10.25080/Majora-92bf1922-00a), SciPy (Virtanen et al., 2020), PyTorch (Paszke et al., 2019), Keras (https://keras.io), statsmodels (https://doi.org/10.25080/Majora-92bf1922-011), scikit-learn (Pedregosa et al., 2011), pybigtools (Huey and Abdennur, 2024), and pyBigWig (Ramírez et al., 2016).

#### Hardware requirements

RepliCNN was developed and tested on a Linux Ubuntu 24.04.1 x86_64 high performance computing (HPC) cluster using two NVIDIA L40s GPUs with CUDA version 12.9 as well as two AMD EPYC 9554 CPU @ 3.1 GHz.

## Availability of data and materials

The human TrAEL-seq data generated in this study is deposited on the Gene Expression Omnibus (GEO) and will be made available upon publication. The public TrAEL-seq, OK-Seq, and GLOE-Seq datasets used in this study are freely available on the Gene Expression Omnibus (GEO) under accession numbers GSE154811 (TrAEL-seq yeast), GSE33786 (OK-Seq yeast), and GSE134225 (GLOE-Seq yeast and human). The high-resolution Repli-Seq data for human was downloaded from GSE137764, and yeast replication timing was inferred from GSE36045.

## Competing interests

The authors declare that they have no competing interests.

## Funding

This works was funded by the Deutsche Forschungsgemeinschaft (DFG, German Research Foundation) via project ID 393547839 – SFB 1361, to K.Z., H.D.U., V.R. and M.C.C., via project ID 533767322 – EXC 3113/1, Cluster for Nucleic Acid Sciences and Technologies – NUCLEATE, to K.Z., and via project ID 529989072 – CA 198/20-1, to M.C.C. We gratefully acknowledge the IMB Genomics Core Facility and its NextSeq 2000 sequencer (funded by the DFG – INST 247/870-1 FUGG).

## Author contributions

D.S. developed RepliCNN, performed all bioinformatic analyses, and wrote the first draft of the manuscript. N.Z. and M.K.P. performed the TrAEL-seq experiments. D.S. and K.Z. designed the study and revised the manuscript with input from all authors. K.Z., V.R., H.D.U., and M.C.C. contributed funding, resources, and guidance.

## Acknowledgments

We would like to express our gratitude to the Genomics and Bioinformatics Core Facilities of the IMB gGmbH (Mainz, Germany) for their assistance in sequencing and data processing. We thank Maximilian Reuter, Mario Keller and all members of the Zarnack group for helpful discussions.

## Notes

### Competing Interest Statement

The authors have declared no competing interest.

